# Regulatory architecture and standing variation drive parallelism in floral evolution

**DOI:** 10.1101/2025.10.28.684842

**Authors:** Kevin Sartori, Natalia Wozniak, Anahid Powell, Ushio Fujikura, Christian Kappel, Tian-Feng Lü, Yang Dong, Stefanie Rosa, Michael Lenhard, Adrien Sicard

## Abstract

Evolution repeatedly gives rise to similar phenotypes, suggesting that constraints and biases make some evolutionary trajectories more likely than others, yet the mechanisms underlying this predictability remain poorly understood. Using repeated transitions to self-fertilization in a model Brassicaceae genus, we show that independent reductions in petal size arise from dosage changes at the same pleiotropic growth regulator, *JAGGED* (*JAG*). Petal development exhibits particularly high sensitivity to *JAG* dosage, resulting in organ-specific morphological shifts with minimal effects on other organs and limited pleiotropy. In both selfing species, these changes were drawn from functionally divergent variants whose polymorphism patterns are consistent with long-term maintenance under weak selection. Our results demonstrate how developmental network architecture and the maintenance of functional variation at pleiotropic loci combine to channel evolution toward predictable outcomes.

## Main

Evolution often produces similar traits across unrelated lineages, suggesting that outcomes are shaped by predictable constraints rather than chance. Such convergent phenotypes often arise in species exposed to similar ecological pressures, implying that adaptive solutions are limited ^1^. Beyond ecology, developmental constraints within Gene Regulatory Networks (GRNs) also restrict the range of viable phenotypes ^2,3^. Because GRNs carry historical legacies, closely related species often share the same constraints and thus respond similarly to shifts in selection pressures ^4^. Although specific cases of genetic convergence have been identified, the general mechanisms that drive repeated phenotypic evolution remain poorly understood ^5^.

Cross-pollination is common in plants and often enforced by self-incompatibility systems, which make reproduction dependent on animal pollinators ^6–10^. To attract pollinators, plants invest in large floral displays ^11–13^. Yet self-incompatibility has repeatedly broken down throughout evolution, producing self-fertilizing (selfing) lineages, likely as reproductive assurance when mates are scarce ^14^. Remarkably, independent transitions to selfing often converge on similar floral changes, collectively termed the selfing syndrome, with a hallmark being a pronounced reduction in flower and petal size ^6,14^. Genetic studies indicate that floral traits often evolve independently from one another or from vegetative traits, suggesting adaptive changes targeting specific organ dimensions ^15–23^. Yet because most growth regulators in plants are pleiotropic, influencing multiple organs ^24^, such specific changes likely depend on rare organ-specific regulators or on regulatory elements within pleiotropic genes. It remains, nevertheless, unclear which general growth regulators can be modulated at an organ-specific level and how such targeted changes in organ size occur. If such a regulation is limited, it could help explain phenotypic convergence following transitions to selfing. Beyond pleiotropy, the effect size and availability of adaptive variation in ancestral populations may further constrain evolution ^25^.

In this study, we exploit the *Capsella* genus as a model to study the convergent evolution of flowers after independent transitions to selfing ^26–33^. In this genus, the selfing species *C. rubella* (*Cr*) and *C. orientalis* (*Co*) evolved independently from an outcrossing *C. grandiflora* (*Cg*)-like ancestor following the breakdown of self-incompatibility, diverging ∼100 kya and within the last 2 million years, respectively (**Fig. 1a**) ^28,34–36^. Both exhibit a fully established selfing syndrome, including a pronounced reduction in petal size (**Fig. 1**) ^37^.

**Fig. 1:**
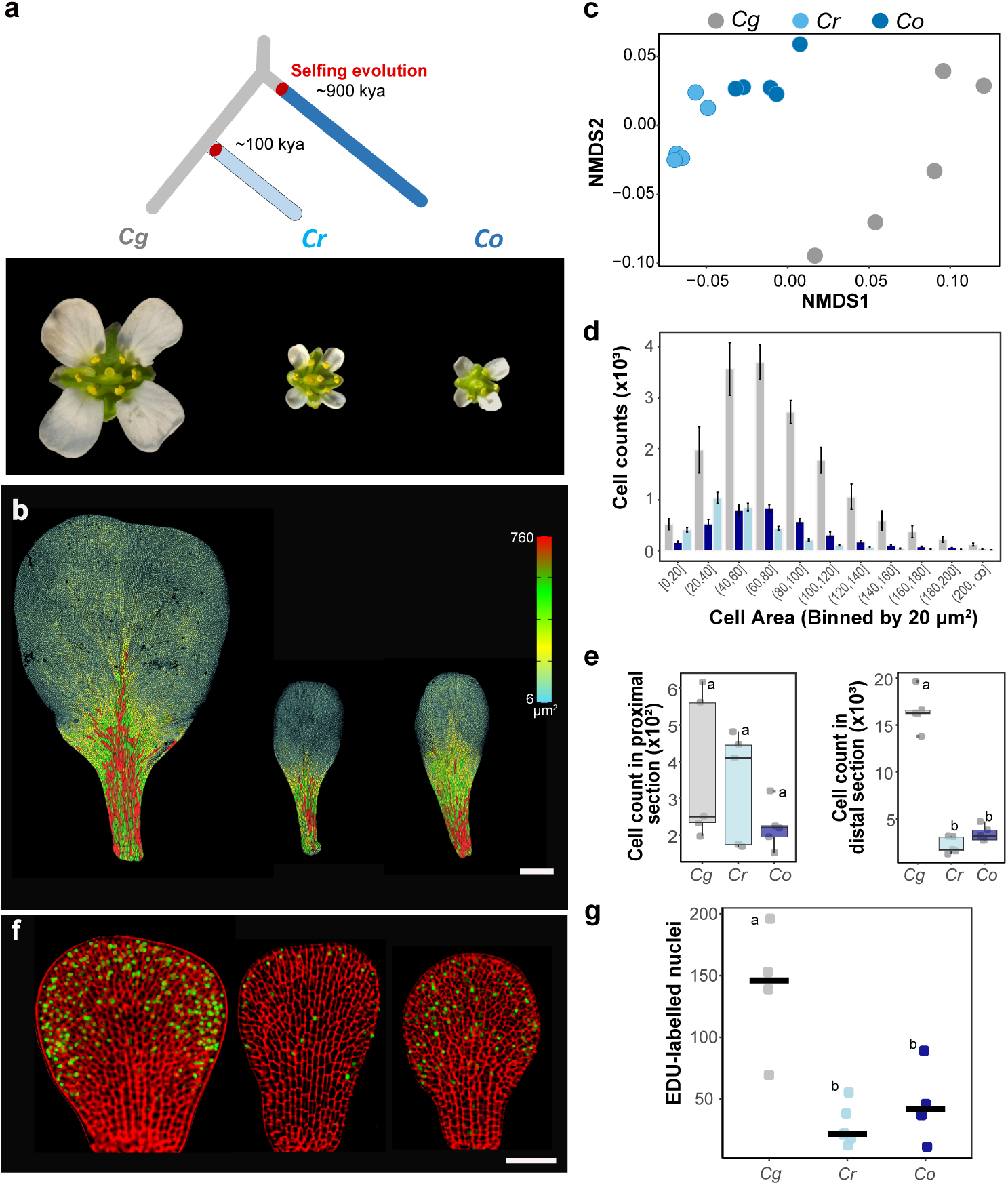
Convergent evolution of petal size following the transition to selfing in *Capsella*. **a**, Schematic representation of the evolutionary history within the *Capsella* genus. **b,** Heatmap illustrating the distribution of cell size within the petals of the three *Capsella* species. Scale bar: 400 μm. **c,** Non-metric multidimensional scaling (NMDS) plot based on Bray-Curtis dissimilarities in cellular features among *C. grandiflora*, *C. rubella*, and *C. orientalis* petals (n = 5 per species). **d,** Average cell count per cell area bin (n = 5 per species). **e,** Cell counts within the proximal and distal regions of the petals. **f,** Representative images showing the distribution of cell division events across the epidermis of stage 1 petals. Fluorescently labelled nucleotide analogue 5-ethynyl-2’-deoxyuridine (EdU; green) was used to visualise newly synthesised DNA, and primary cell walls were stained with propidium iodide (PI; red). Scale bar: 50 μm. **g,** Quantification of EdU-labelled nuclei in stage 1 petals. Letters indicate significant differences as determined by one-way ANOVA with Tukey’s HSD test (α = 0.05).

### Convergent petal size reduction via similar reduction of distal cell proliferation

A major phenotypic change in *Cr* and *Co* is a pronounced reduction in petal size, accompanied by fewer cells, implicating altered cell proliferation as a central mechanism ^38^. To examine the developmental basis of these changes, we analyzed cellular patterns along the proximal-distal axis of petals, incorporating various descriptors of cell shape and directional growth (**Extended Data Table 1**). Petals exhibit two distinct cell clusters. A proximal cluster of large, elongated cells and a distal cluster of small, rounded cells, separated by a transition zone where cell size and anisotropy gradually decrease (**Fig. 1B and Extended Data Fig. 1**). While this pattern is largely conserved across species, cellular features along this axis can distinguish the petals of selfers from those of the outcrosser (one-way ANOVA: *F*(1,13) = 28.54, *p* = 0.00013, *η²* = 0.687) (**Fig. 1b, 1d, and Extended Data Fig. 1, and 2**). Selfers differ primarily in the distal region, where cell number is reduced roughly five-fold compared to the outcrosser, resulting in a two-fold reduction in length and a three-fold decrease in width (**Fig. 1e and Extended Data Fig. 1c; Extended Data Table 2**). Early petal growth occurs via cell proliferation, which initially takes place throughout the primordia. Once the petal primordia reach approximately 90 µm in length, proliferation rapidly becomes restricted to the distal region, thereby determining the size of the distal section, while cell elongation is largely confined to the proximal domain (**Fig. 1 and Extended Data Fig.3**). In selfers, cell proliferation in the distal margin, as judged from the number of S-phase cells, is reduced three- to four-fold relative to the outcrosser, consistently across both lineages (**Fig. 1f and 1g**). Thus, despite independent origins, petal size reduction in these selfing species arises through the same developmental mechanism: restricted growth at the distal margin due to decreased cell division.

### Independent selfing lineages co-opted *JAG* to reduce petal size

The convergence in petal development suggests that similar genetic mechanisms underlie size reduction in the two selfing species. Previous studies indicated that half of the loci associated with petal size are shared between *Cr* and *Co*, with PAQTL2 on chromosome 2 contributing most to phenotypic evolution (**Extended Data Fig. 4a**) ^38,39^. To refine the location of PAQTL2 in each selfing species, we introgressed the outcrossing *Cg* alleles into each selfer, creating near-isogenic lines (*NILrg1* and *NILog2*) and screened for recombinants within this region (**Fig. 2**). By analyzing petal size segregation in the progeny, we narrowed down the causal mutation (s) to 14.6 kb in *Cr* and 130 kb in *Co*. The most informative recombinants from *NILrg1* indicated that the causal mutation(s) are situated between 9,061,779 and 9,076,392 bp on chromosome 2. We confirmed this region by generating a quasi-isogenic line (*qILrg*) for the 14.6-kb interval, where *Cr* homozygotes exhibited ∼20% smaller petals than *Cg* homozygotes, recapitulating the PAQTL2 effect (**Extended Data Fig. 4c and 7a**). In *NILog2*, the most informative recombinant localized the causal mutation to 9,019,066–9,149,619 bp on chromosome 2, accounting for a 25% petal reduction in *Co* (**Fig. 2b and Extended Data Fig. 7a**). This region overlaps with the locus identified in *Cr*, confirming that the same genomic locus underpins PAQTL2 in both selfing species.

**Fig. 2.**
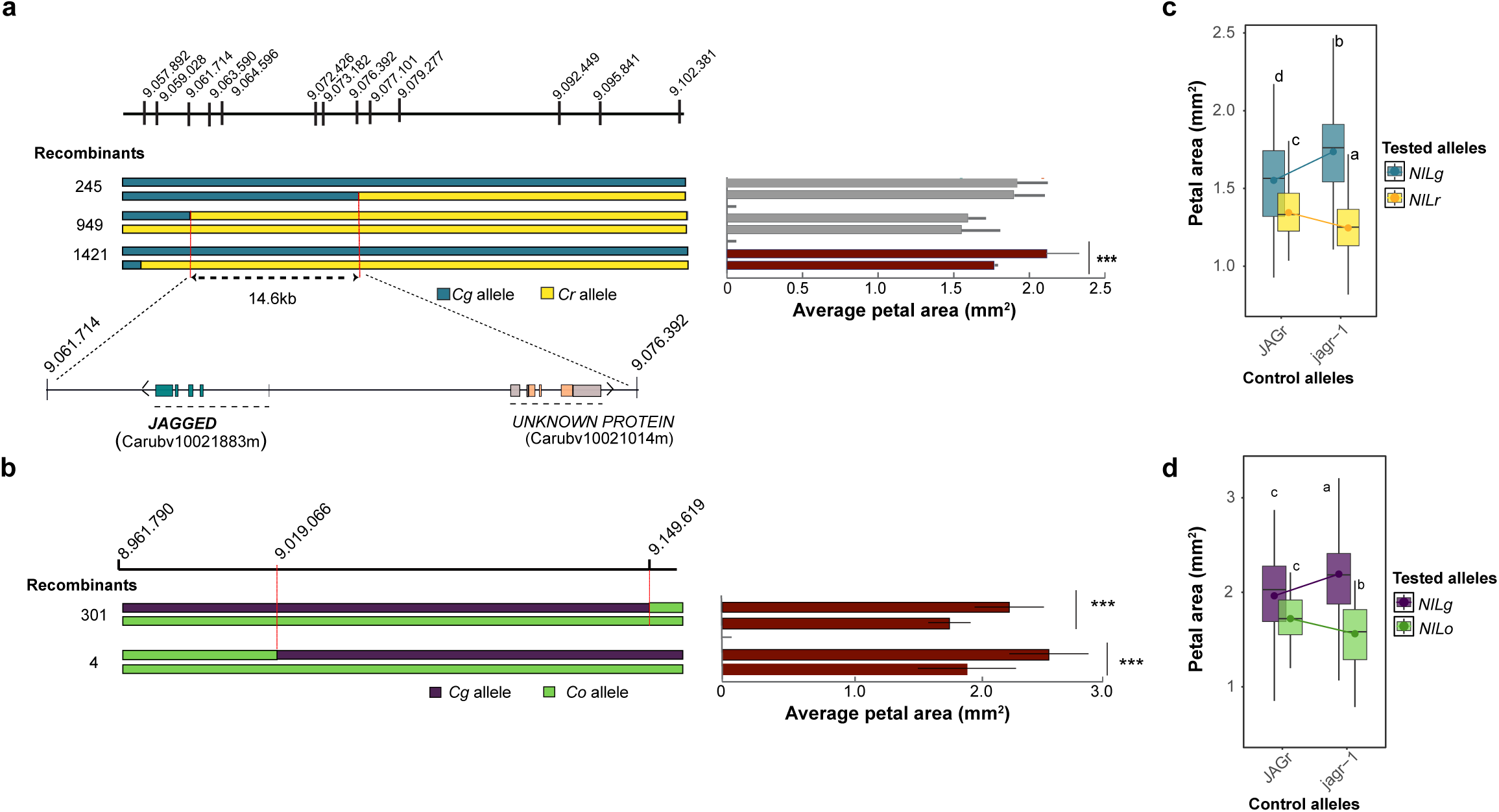
Allelic variation within *JAG* contribute to petal size reduction in indepdent selfing lineages. **a** and **b** Fine mapping of PA_QTL2 in *C. rubella* (*Cr*) and *C.orientalis (Co)*, respectively. The color coded bars on the left repsent the genotype of the most informative recombinants. The position on scaffold 2 are indicated on the top in base pair (bp). The bar chart on the right shows the average petal size for each alternative homozygous plants for each recombinant lines. Red filled bars indicate significant differences between alternative homozygous genotypes, *P* <0.05 (Student’s t-test). **c** and **d** Quantitative complementation assay illustrated with an interaction plot of the crosses between *NILrg* **c** or *NILog* **d** with *jagr-1*+/-. Letters indicate significant differences as determined by one-way ANOVA with Tukey’s HSD test (α = 0.05).

The 14.6-kb interval contained only two genes, including the *Arabidopsis thaliana JAGGED* (*JAG*) ortholog, a zinc-finger transcription factor, known to control lateral organ growth and distal cell proliferation (**Fig. 2a**) ^40^. In an EMS-mutagenized *C. rubella* screen, we identified a mutant phenocopying the *A. thaliana jag* mutation (**Extended Data Fig. 6a**)^40^. Bulked-segregant analysis revealed a splice-site mutation in the third exon of *JAG*, preventing proper splicing and introducing a premature stop codon (**Extended Data Fig. 5**). The defect was rescued by reintroducing a wild-type *Cr* allele, confirming *JAG* inactivation as causal (**Extended Data Fig. 6b**). We thus used this mutant in a quantitative complementation test ^41^ to test whether *JAG* underlies PAQTL2 in both selfers. In this assay, natural alleles were crossed to a jag+/– background, where functional differences between the natural alleles are expected to be amplified only when the loss-of-function mutation affects the same gene. Consistently, in both cases, the differences between the selfer and outcrosser alleles were stronger when combined with the mutant alleles (**Fig. 2c and d**). Linear model analysis revealed a significant interaction between the tested (*Cg1* and *Cr* or *Cg2* and *Co*) and control allele (WT and *crjag*) (*β* = −0.38976 ± 0.08710, *t* = −4.475, *p* = 8.56 × 10⁻^6^ for *NILog*; and *β* = 0.311 ± 0.043, *t* = 7.245, *p* = 2.27 × 10⁻^12^ for *qILrg*), indicating that polymorphism within *JAG* contributed to a reduction in petal size in both cases. To further confirm *JAG’s* role, we transformed genomic constructs corresponding to the *Cg* (*JAGg1*) and *Cr* (*JAGr*) alleles segregating in *NILrg*, as well as the *Cg* (*JAGg2*) and *Co* (*JAGo*) alleles from *NILog*, into the *A. thaliana jag-2* mutant (**Extended Data Fig. 4d and e**). All alleles rescued the mutant phenotype, but *Cg1* and *Cg2* alleles produced larger petals than *Cr* and *Co* alleles, respectively. These results further support that allelic variation in *JAG* underlies convergent petal size reduction in both selfing species.

### *JAG* is essential to promote distal petal cell proliferation

Inactivation of *JAG* in *Cr* causes pleiotropic effects, leading to a ≥10% reduction in the size of most organs, with particularly strong effects on leaves (30%), petals (60%) (**Extended Data Fig. 6**), and fruit ^42^. To uncover the developmental basis, we compared wild-type and *crjag* petals. Silhouette overlays showed that the proximal petal region was largely unaffected, whereas the distal region was almost entirely missing (**Fig. 3a and 3b**). Cellular analysis revealed a ∼5-fold reduction in total cell number (**Fig. 3c and 3d**). This reduction in cell number was particularly pronounced in the transition and distal zones (**Fig. 3d**). Cell elongation in the proximal region and the position of the transition zone were largely unaffected, indicating that overall petal growth patterning is not altered by *JAG* (**Fig. 3b**). However, growth in the marginal part of the petal was almost completely abolished, with only a few small rounded cells formed within the distal area. This was associated with more than a twofold reduction in the number of cell proliferation events at the onset of cell proliferation within the distal part of the petal primordia (**Fig. 3e and f**). Together, these results indicate that *JAG* is essential for promoting and maintaining a high rate of cell proliferation at the petal margin. Lack of function almost completely abolishes growth in the distal region. These observations suggest that *JAG* does not affect the overall growth pattern of the petal but instead promotes cell proliferation within the distal margin, providing strong support for its proposed function as a distal growth promoter during petal development ^43^ and explaining the cellular basis of convergent petal size reductions in *Capsella* (**Fig. 1**).

**Fig. 3.**
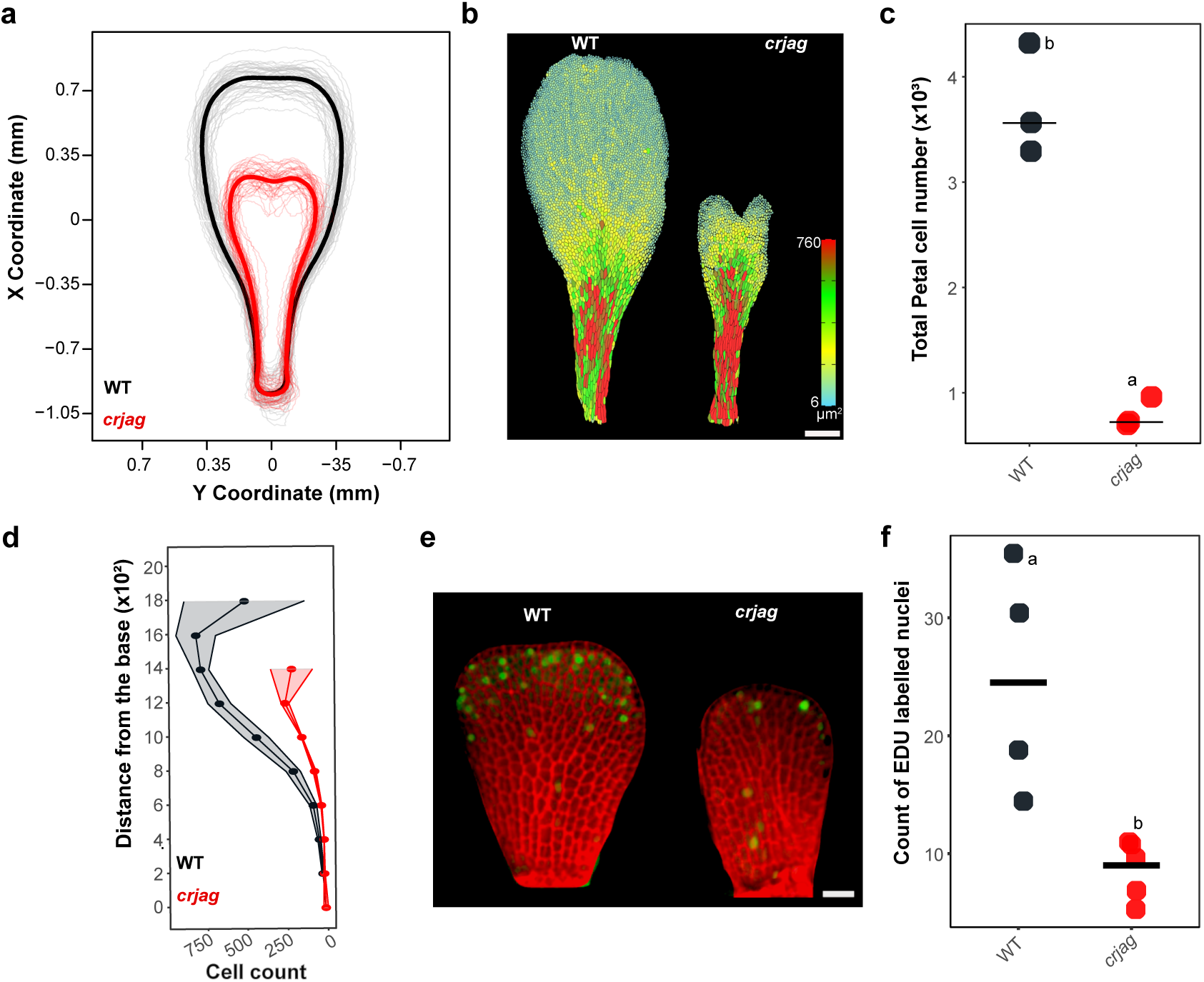
Loss of *JAG* function strongly inhibits cell proliferation during petal growth. **a**, Average shape of *Cr* and *crjag* petals. **b,** Heatmap illustrating the distribution of cell size within the petals of *Cr* and *crjag*. Scale bar: 200 μm. **c,** Total number of cell within *Cr* and *crjag* petals (n = 3 per genotype). **d,** Average cell count per 200 μm section along the longitudinal petal axis (n = 3 per genotype). Numbers on the y-axis indicate the starting point of each section. **e,** Representative images showing the distribution of cell division events across the epidermis of stage 1 petals. Fluorescently labelled nucleotide analogue 5-ethynyl-2’-deoxyuridine (EdU; green) was used to visualize newly synthesized DNA, and primary cell walls were stained with propidium iodide (PI; red). Scale bar: 20 μm. **f,** Quantification of EdU-labelled nuclei in stage 1 petals. Letters indicate significant differences as determined by one-way ANOVA with Tukey’s HSD test (α = 0.05).

### Cis-regulatory changes in *JAG* drive petal size reduction in selfing *Capsella* species

To assess the impact of allelic variation at *JAG*, we compared petal morphology in the *NILog* and *qILrg* lines (**Fig. 4 a-d**). Consistent with the loss of *JAG* function, both showed a selective reduction in distal petal growth, with total petal area decreased by 22% and 15%, respectively (**Extended Data Fig. 7**). This reduction reflected a lower total cell number, most pronounced at the distal tip. Strikingly, the effect was specific to petals; other floral organs and leaves were unaffected, in contrast to the broad reductions seen in *crjag*. These results suggest that *JAG* activity has been selectively modified during petal development.

**Fig. 4.**
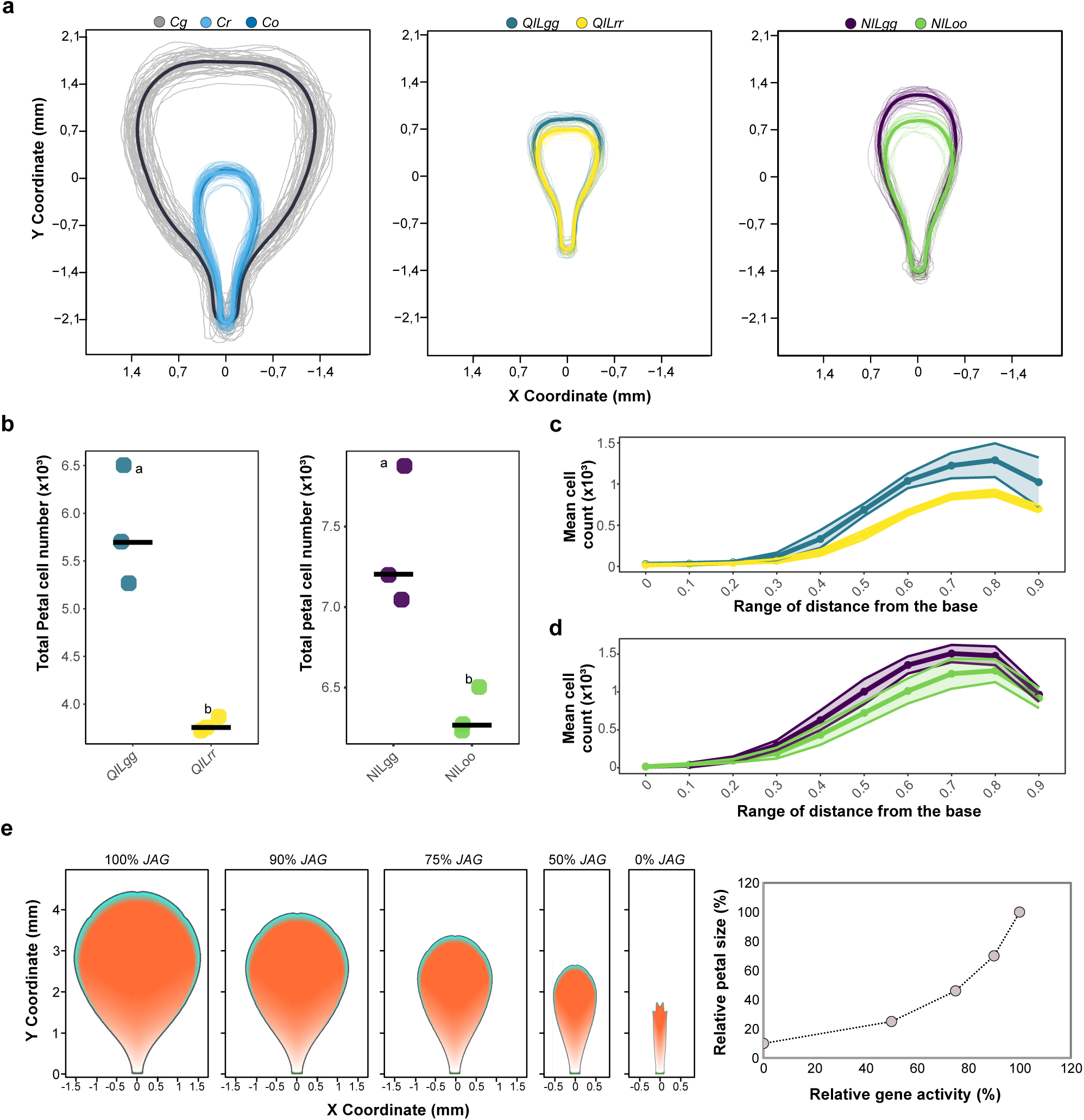
Developmental constraint on petal size evolution via heightened *JAG* sensitivity. **a**, Average petal shape of *Cg*, *Cr* and *Co* petals (left panel), of quasi Isogenic lines (*QIL*) homozygous for the *Cg1JAG* (QILgg) or *CrJAG* (QILrr) allele (middle panel) and of Near Isogenic lines (*QIL*) homozygous for the *Cg2JAG* (*NILLgg*) or *CoJAG* (*NILLoo*) allele (right panel). **b,** Total number of cell within *QILgg* and *QILrr* and within *NILgg* and *NILoo* petals (n = 3 per genotype). **c-d**, Average cell count for 10 section along the longitudinal axis within *QILgg* and *QILrr* (**c**) and within *NILgg* and *NILoo* (**d**) petals (n = 3 per genotype). Numbers on the y-axis indicate the starting point of each section. **e,** Output of the *Capsella* petal development model with a divergent polarity field. The effect of *JAG* on basic growth parameters was progressively reduced, as indicated at the top of each graph from left to right. The rightmost graph shows the impact of decreased *JAG* activity on petal size. Letters indicate significant differences as determined by one-way ANOVA with Tukey’s HSD test (α = 0.05).

Among the polymorphisms distinguishing the tested *JAG* alleles, four resulted in amino acid substitutions, but none affected conserved regions or the C2H2 zinc-finger domain. Furthermore, promoter-swap experiments revealed a cis-regulatory basis. *A. thaliana jag-2* mutants transformed with the *Cr* coding sequence under the *Cg1* promoter (*pJAGg1::JAGr*) developed significantly larger petals than those carrying the reciprocal construct (*pJAGr::JAGg1*) (Fig. S4 D). Moreover, *jag-2*; *pJAGr::JAGg1* petals were similar in size to *jag-2*; *JAGr* petals and significantly smaller than *jag-2*; *JAGg1* petals (**Extended Data Fig. 4**). Thus, in *Cr*, differences in *JAG* activity are explained by cis-regulatory rather than protein-coding changes. Because both selfers share the same non-synonymous substitutions, these are unlikely to contribute in Co, also suggesting cis-regulatory evolution in this species.

### Subtle changes in *JAG* dosage drive petal size evolution with minimal pleiotropy

Single-cell analysis revealed reduced *JAG* expression in both developing leaves and petals of the selfers compared to the outcrosser (**Extended Data Fig. 8**). Because *JAG* promotes cell division ^44^, its reduced activity likely explains the lower proliferation observed at the petal margin. Yet, only petals showed strong size reduction, suggesting heightened sensitivity to *JAG* dosage. This is consistent with the severe petal defects of *jag* mutants and the established role of *JAG* in driving distal petal growth.

To test whether *JAG* dosage could explain petal size differences, we used an existing petal growth model based on the Growing Polarized Tissue (GPT) framework ^45^ (**Extended Data Fig. 9**). In this model, tissue polarity is generated by diffusion of a polariser (POL) produced by the basal organiser (PROXORG), which drives growth along the proximal-distal axis (Kpar). The distal factors (DGRAD, DISTORG) promote lateral (Kper) and divergent growth from the petal midline, respectively. The model reproduces *A. thaliana* petals and is consistent with *JAG* functioning as DGRAD, with the phytohormone auxin signalling contributing to the polarity field establishment ^45^. However, it failed to capture *Capsella* petal shapes (**Extended Data Fig. 9**). Adjusting the model by broadening DGRAD and DISTORG domains and allowing DGRAD to also influence Kpar yielded realistic *Cg* petals (**Fig. 4, Extended Data Fig. 9 and Methods**). With this new model, the complete loss of DGRAD, combined with fragmented auxin distribution (as in ^45^), produced short and narrow ‘jagged’ petals phenocopying the *crjag* mutants. Quantitative tuning showed that even modest reductions in *JAG* activity had strong effects on petal size. A 10% decrease in its effects on Kper and Kpar led to a 23% reduction in petal size, while a 25% reduction in *JAG* resulted in more than a twofold decrease in petal size (**Fig. 4e**).

To test whether organ-specific sensitivity could explain the minimal pleiotropy of change in *JAG* expression, we next used an existing leaf growth model ^46^ (**Extended Data Fig. 10**). In this framework, a PGRAD gradient along the proximal-distal axis regulates growth parallel to the axis (Kpar), while the broadly expressed factor LAM promotes perpendicular growth (Kper), which is further inhibited by MID along the central axis. At later stages, LATE reduces Kpar and enhances Kper activation by LAM. Because *crjag* leaves show a 14% reduction in length and 18% in width, we hypothesised that *JAG* contributes to both PGRAD and LAM activity, consistent with its expression at proximal side of leaf and sepal margins ^40,47,48^. Calibrating the model to match the *crjag* phenotype and running both the petal and leaf models for a similar number of iterations under different *JAG* dosage revealed that petals were far more sensitive to *JAG* reduction. At 75% activity, petal area had already decreased by ∼30%, whereas leaf area was reduced by less than 15%, and this contrast becomes even stronger at lower dosage levels (**Extended Data Fig. 10**). This analysis shows that modest decreases in *JAG* activity strongly affect petal growth but have only minor effects on other organs. Such heightened sensitivity may represent a developmental constraint and help explain why *JAG* has been repeatedly co-opted to reduce petal size in different lineages. Because *JAG* plays a central role in promoting petal growth while contributing less to other organs, even subtle changes in its activity can drive major evolutionary shifts in petal morphology with minimal pleiotropic consequences.

### Standing variation facilitates parallel petal size evolution

A notable observation was that polymorphisms affecting *JAG* function in both selfing lineages showed similar overdominant effects (**Fig. 5**). In both *NILog* and *qILrg* heterozygotes had significantly larger petals than homozygotes (effect sizes of 0.342 and 0.186 mm²; SE = 0.132 and 0.0871; df = 19 and 20; t = 2.582 and 2.136; Bonferroni-adjusted *p* = 0.0183 and 0.0452, respectively). Similarly, in complementation crosses, *Cg/crjag* heterozygotes produced larger petals than *Cg/Cr* heterozygotes (**Fig. 2**). *JAG*, therefore, appears to be prone to overdominant mutations, likely due to dosage-sensitive responses in petal development.

**Fig. 5.**
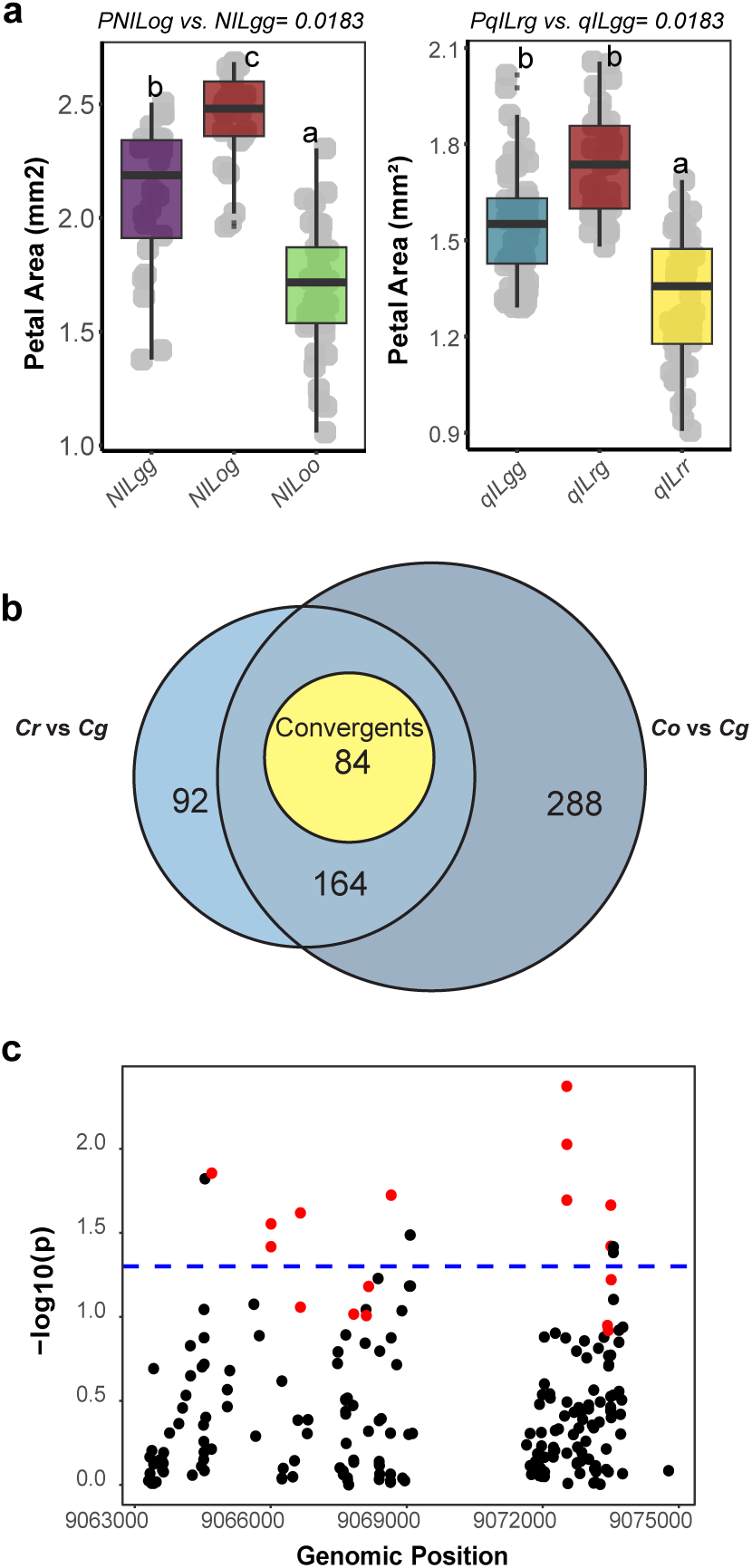
Standing genetic variation in Cg contribute to convergence. **a**, Relationship between petal size and *JAG* genotypes in the *NILog* and *qILrg* lines. Data points are gray dots; boxplots show petal area distribution per genotype. Letters denote significant differences from Tukey tests after a mixed-effects model with plant as a random effect. Bonferroni-adjusted p-values compare heterozygotes to larger homozygotes. **b,** Venn diagram depicts lineage-specific and shared polymorphic sites between the two Capsella selfer lineages versus their outcrosser ancestor. Convergent polymorphisms, where the same allele was fixed in both of the selfers, are highlighted in yellow. **c,** Manhattan-like plot displays genotype-phenotype (petal length) associations and overdominant loci. Each point tests a genomic locus’s association with petal length; the x-axis shows position; the y-axis shows -log10(p), significance from linear model testing genotype effect on the phenotype. The dashed line marks p=0.05. Overdominant loci, where heterozygotes show the largest mean, are in red.

Such overdominance, if maintained by selection, could promote allele reuse by keeping functionally divergent variants in the outcrosser population. To test this, we examined allele frequencies across natural populations at polymorphic sites segregating between the haplotypes used in transgenic assays (*Cg1-Cr* and *Cg2* - *Co*). Of 642 polymorphic sites, 84 converged between selfers. Yet, none of these sites were fixed, as selfer alleles still occurred at ≥4% in *Cg*. This lack of strong differentiation excludes a scenario of parallel evolution via identical independent changes at the same locus post-divergence. Both selfers also carried species-specific alleles (16 in *Cr*, 26 in *Co*), indicating that independent de novo changes may also have contributed (**Extended Data Fig. 11**). Because the high number of convergent polymorphisms suggested evolution via standing genetic variation, we tested whether selfer/outcrosser polymorphisms segregating in *Cg* were associated with petal size (**Fig. 5 and Extended Data Fig. 12**). Among 233 polymorphisms with sufficient genotype counts, 15 were significantly correlated with petal length (FDR-adjusted P < 0.05). Of these, nine showed overdominance, with heterozygotes having larger petals. Six of these nine were fixed in both selfers, and five were associated with smaller petals (**Extended Data Fig. 12**). This suggests that collateral evolution through the fixation of multiple small-effect variants could also contributed to petal size reduction.

Because *Cg/Cr* and *Co* lineages diverged approximately 1 Mya, this scenario implies that these polymorphisms have persisted for thousands of generations (**Fig. 1**, ^49,50^). Such persistence could plausibly be explained by heterozygote advantage, given the importance of petal size for pollinator attraction. However, none of the convergent overdominant polymorphisms showed significant deviations from Hardy-Weinberg equilibrium (HWE) or evidence of heterozygote excess in *Cg* (**Extended Data Fig. 13**). Nonetheless, in a population with a large effective population size, such as *Cg* ^51^, weak overdominant selection could maintain allele frequencies near a stable equilibrium without causing apparent departure from HWE ^52^. To test this hypothesis, we estimated selection coefficients acting on petal morphology by correlating petal size with reproductive efficiency in a hybrid outcrossing population (**Extended Data Fig. 13b**). Using these estimates, together with the observed heterozygote effects, we modelled allele trajectories for each convergent overdominant polymorphism (**Extended Data Fig. 13c**). Despite low selection coefficients (average *s* against homozygotes ≈ 0.0056), overdominant selection was sufficient to maintain these polymorphisms over millions of generations (probability of heterozygote maintenance: 1 with selection and drift vs. 0.18 under drift alone) (**Extended Data Fig. 13c**). Consistent with our above analysis, only rare simulations (<0.05%) led to HWE departures. Together, these results indicate that weak overdominant selection can maintain functionally divergent alleles at polymorphic frequencies in large outcrossing populations, thereby enabling repeated recruitment of standing variation and facilitating phenotypic convergence through collateral evolution.

### Implications for convergent evolution of selfing lineages

Petal reduction in both *Capsella* selfers arose through the identical developmental mechanism and the co-option of the same genetic solution (**Extended Data Fig. 14**). In both lineages, cis-regulatory changes reduce the activity of *JAG*, a general growth promoter, leading to decreased cell proliferation in the distal petal margin. Although reduced *JAG* expression is detectable in several organs, only petals are strongly affected, likely because petal growth is more sensitive to *JAG* dosage. This heightened sensitivity appears to impose a developmental constraint on evolution by enabling large morphological changes in petals while limiting pleiotropic effects, explaining at least in part gene reuse and the convergent evolution observed between independent transitions to self-fertilisation. In addition, the reuse of standing variation also appears to have facilitated phenotypic parallelism. Variants segregating at intermediate frequency in *Cg*, where they influence petal size, have been similarly fixed in both selfing lineages. Weak overdominant selection associated with improved pollinator attraction explains the long-term persistence of these variants in *Cg,* and thus their repeated use. Together, our findings illustrate how the interplay between standing genetic variation and developmental constraints interact to bias evolutionary trajectory toward repeatable outcomes.

## Methods

### Plant material

This study used *Capsella rubella*, *C. orientalis*, and *C. grandiflora* accessions previously described by ^37^. Near-isogenic lines (NILs) segregating for PAQTL_2 were generated by introgressing the *Cg926* allele into *Cr1504* (*NILrg*) and *Co1983* (*NILgo*) through five and three rounds of backcrossing, respectively, followed by two additional generations of selfing while maintaining heterozygosity at the PAQTL_2 locus. Progeny from these two NILs were used for fine mapping. The crossing scheme used to generate the quasi-isogenic line (qILrg) is shown in **Extended Data Fig. 4c**.

The *jag-2* EMS mutant in *Landsberg-erecta* (L*-er*) background showing recessive floral phenotype was previously identified to have a single nucleotide change (C/T) in the second exon resulting in a premature stop codon ^40^. Here, we have identified the *Crjag* mutant in *C. rubella Cr22.5* background by screening a *C. rubella* (*Cr22.5*) ethyl-methanesulfonate (EMS) mutagenized population for plants with reduced petal size ^53^. The genetic basis and phenotypic characterization of this mutant are presented in this manuscript.

### Growth conditions

Seeds were surface-sterilized, plated on half-strength Murashige-Skoog medium supplemented with gibberellic acid (Duchefa Biochemie; 3.3 mg/L GA), and vernalized for two days at 4°C. Germination occurred at 22°C for ten days. To initiate the transition to flowering, seedlings were placed at 4°C for ten days. Subsequently, plants were grown in a growth chamber under long-day conditions (16 hours light/8 hours dark) with a temperature cycle of 22°C during the day and 16°C at night, 70% humidity, and a light intensity of 150 μmol m⁻²s⁻¹.

### Morphological measurements

To characterize the effect of PAQTL_2, diverse genetic constructs, and *jag* mutations, sizes of the various organs were measured. Sepals, petals and anthers were measured from the 15th and 16th fully opened flowers on the main stem. Carpel length was measured from unfertilized flowers. Six petals, six sepals, six anthers and two carpels were measured per plant. Dissected flower organs were flattened and scanned at a resolution of 3200 dpi (HP ScanJet 4370). Measurements were conducted on digitized images by using ImageJ ^54^. Leaf size was measured on the fully expanded 6th leaf. The leaves were flattened and scanned at a resolution of 300 dpi using an HP ScanJet 4370. Measurements were then taken from the digitized images using ImageJ. ^54^.

### Morphometric analysis of petal outlines

Morphometric analysis of petal shape described as closed outlines was performed using Elliptic Fourier Descriptors (EFD) ^55^. First, the digital images were converted into binary images using ImageJ ^54^. Coordinates of boundaries were extracted using the Python module scikit-image ^56^. Outline coordinates were then analysed and visualised using the Momocs package implemented in R ^57^.

### Developmental analysis of petal growth

Petal cell size and cell number were determined from dried-gel agarose prints ^58^ of whole petals from fully open flowers. Cell outlines were captured using and Olympus BX 51 light microscope and AxioCam HRc camera (Carl Zeiss). The cell-outline images were merged using the photomerge function of Adobe Photoshop and further processed with the Python module scikit-image. First, cell-outlines were segmented using adaptive thresholding. Then, binary images of cell borders were dilated, skeletonized and curated by overlapping them with the original images in GIMP (https://www.gimp.org/). Cell areas and centroid coordinates were extracted and used to estimate the average cell number and area per segment along the petal longitudinal axis. The analysis of differences in cell number and size between genotypes was performed using R ^59^. Curated segmented images were then analysed with ImageJ by defining Region Of Interest (ROI) and measuring diverse morphological features for each particle. Heatmaps were generated in ImageJ using the ROI colour coder function.

### Confocal imaging

To visualize petal primordia, young flower buds were fixed in a solution containing 50% methanol and 10% acetic acid. mPS-PI staining was carried out following the protocol by Ichihashi et al. (2011)^60^, with minor modifications. Imaging was performed using a confocal laser scanning microscope (LSM780, Carl Zeiss), with excitation at 488 nm and emission collected between 520 nm and 720 nm. Z-stack images were acquired at a resolution of 512 × 512 with a step size of 0.5–1 µm along the z-axis. Image stacks were processed and analyzed using MorphoGraphX ^61,62^. Surfaces were detected using the “edge detect” tool with a threshold of 10,000. Initial meshes were generated with 5 µm cubes and subsequently subdivided three times before projecting the mPS-PI signal located 4-6 µm from the mesh surface.

### EDU staining

Cell division events were visualized using a combination of 5-ethynyl-2’-deoxyuridine (EdU) incorporation and modified pseudo-Schiff propidium iodide (PI; Sigma-Aldrich) staining, following the protocol described in Riglet et al. (2024) with minor modifications ^63^. Inflorescences were harvested and placed on EdU staining medium consisting of 0.22% (w/v) Murashige and Skoog basal salts with vitamins (Duchefa Biochemie, M0222.0050), 3.5% (w/v) sucrose, 0.8% (w/v) agarose, and 10 μM EdU (Invitrogen A10044, Thermo Fisher Scientific), adjusted to pH 5.7. Samples were submerged in the same medium lacking agarose and incubated for 15 hours in a growth chamber under long-day conditions (16-hour light/8-hour dark photoperiod) at 22°C, with an average light intensity of 150 μmol m⁻²s⁻¹. Following incubation, samples were dehydrated through a graded ethanol series (25%, 50%, 75%, 90%, and 100%; 10 minutes each), then rehydrated through the same series in reverse. Tissues were then treated overnight at 37°C with 20 mM α-amylase (Sigma-Aldrich A4551) in phosphate buffer (pH 7.0) supplemented with 2 mM NaCl and 0.25 mM CaCl₂. After washing twice with water and once with 100 mM Tris buffer (pH 8), the samples were incubated for 1 hour in a solution containing 10 μM Alexa Fluor 488-azide (Invitrogen A10266) in 100 mM Tris buffer (pH 8). Subsequently, tissues were incubated for 30 minutes in a click-reaction solution containing 10 μM Alexa Fluor 488-azide, 100 mM Tris (pH 8), 1 mM CuSO₄, and 100 mM ascorbic acid. Samples were washed three times with water. Young flower buds were dissected in water to expose petals and incubated in 1% periodic acid for 30 minutes, followed by two water washes. Samples were then stained with 0.01 μg μl⁻¹ PI for 3 hours, cleared for 2 hours in chloral hydrate solution, and mounted in Hoyer’s solution (30 g gum arabic, 200 g chloral hydrate, 20 g glycerol, and 50 ml water). Imaging was performed using a confocal laser scanning microscope (LSM780, Carl Zeiss) with excitation at 488 nm. Emission was collected between 493–547 nm for EdU and 622–718 nm for PI. Z-stack images were acquired at 512 × 512 resolution with a step size of 0.5–1 µm along the z-axis. EdU signals were detected and quantified using ImageJ.

### Genetic mapping

To refine the position of PAQTL_2, we conducted a segregation analysis in the *NILrg* background. Initial mapping using approximately 400 progeny plants identified a 138.095 kb region on chromosome 2, located between markers Bs5 (position 8,964,289) and SH3_2 (position 9,102,384) (**Extended Data Table 3**). To further narrow down this interval, we screened about 1,000 additional progeny plants from NILrg to identify individuals with recombinant chromosomes between these two markers. The positions of the recombination breakpoints were determined by genotyping these recombinants with additional markers within the interval. Selfed progenies from the 15 most informative recombinant lines were then genotyped to identify individuals homozygous for either parental allele. Petal size was quantified in these homozygous plants, and a t-test was used to compare the average petal size between genotype classes for each recombinant to localize the causal mutation in relation to the position of each recombination breakpoint.

For fine-mapping PAQTL_2 in the *NILgo* background, we first identified a *NILgo* line segregating for a ∼1 Mb region surrounding PAQTL_2, located between markers JAGI (position 8,127,464) and m400 (position 9,149,587) on chromosome 2 (**Extended Data Table 4**). Approximately 1,400 *NILgo* progeny plants were screened to identify recombinants between these markers. Each recombinant was genotyped with additional markers within the focal region to pinpoint the recombination breakpoints. Selected recombinant plants were selfed, and progeny were genotyped to identify individuals homozygous for either the *C. grandiflora* or *C. orientalis* allele in the segregating region. As described above, petal sizes of both genotype classes were measured and compared using a t-test, enabling us to further refine the position of the causal mutation.

### Molecular cloning and plant transformation

*C. rubella* (*JAGr*), *C. orientalis* (*JAGo*), two *C. grandiflora* (*JAGg1* and *JAGg2*) and two promoter swap genomic constructs: *pJAGg1::JAGr* and *pJAGr::JAGg1* were used to test the existence of allelic variation influencing petal size within the *JAG* locus. The 8.3 kb *JAGr*, 9.4 kb *JAGg*, 9.4 kb *pJAGg::JAGr* and 8.3kb *pJAGr::JAGg1* genomic constructs were amplified in two PCR reactions from genomic DNA of plants homozygous for the *C. rubella* allele (*NILrr)* and plants homozygous for the *C. grandiflora* allele (*NILgg1)* as described in **Extended Data Table 5**. The 8.3 kb *JAGo* and a 10.1 kb *JAGg2* genomic constructs were amplified in three PCR reactions from genomic DNA of plants homozygous for the *C. orientalis* allele (*NILoo)* and plants homozygous for the *C. grandiflora* allele (*NILgg2)* as described in **Extended Data Table 5**. Primers used are presented in **Extended Data Table 6**. The genomic fragments were first assembled and subcloned into a modified version of pBluescript II KS (StrataGene, pBlueMLAPUCAP) using the InFusion® HD Cloning Plus kit (Clontech) and sequenced. The reconstituted cassettes were transferred into the AscI site of the plant transformation vector pBarMAP, a derivative of pGPTVBAR ^64^. These genomic constructs were then used to transform *Crjag* and *jag-2* plants by floral dip ^65^. Note that since *NILrg* and *NILog* were generated from crosses with different *Cg926* individuals and since *C. grandiflora* is a highly heterozygous outcrosser, the two *C. grandiflora* alleles were not identical in sequence and were therefore named *JAGg1* and *JAGg2*.

### Identification of the *Crjag* mutation

To identify the *Crjag* mutation, we followed a mapping-by-sequencing approach ^66^. 500 F2 progeny of a cross between a mutant that we initially termed *heart-stopping (hts)* because of the defect in fruit development and *Cr1504* were screened to isolate plants with a WT or mutant phenotype. About 100 plants for each genotype were pooled and sequenced using the Illumina NextSeq 500 instruments (2 x 100 cycles). DNA-seq data were processed using Trimmomatic ^67^ to remove adapter sequences. Quality control was done using FastQC (http://www.bioinformatics.bbsrc.ac.uk/projects/fastqc). Reads were then mapped at per sample level against the *C.rubella* reference genome (Cru_183 from phytozome.org) using BWA-MEM ^68^. Variant calling was performed using SAMtools ^69^. Those calls were then further processed using R ^59^ to extract *Cr22.5* (background of *crjag* mutation)/*Cr1504* polymorphisms. The ratio between the number of homozygous and heterozygous calls within 200 kb windows was used to localize the mutation.

### RNA splicing analysis

To detect splicing differences, RNA was first extracted from *Crjag* and *Cr22.5* inflorescences (containing the inflorescence meristem and the flower buds from stage 1 to 9) using Trizol (Life Technologies). RNAs were treated with TurboDNAse (Ambion) and reverse transcribed using oligo(dT) and the Superscript III Reverse Transcriptase (Invitrogen). Primers onw545 and onw546 were used to amplify full-length *JAG* transcripts.

### Modelling of petal growth

The *Capsella* petal growth model is based on a 2D Arabidopsis thaliana petal growth model developed by Susanna Sauret-Güeto *et al.*, 2013 ^45^. Models were specified and manipulated using the MATLAB-based application *GFTbox* ^70^. Growth patterns are guided by spatially distributed factors across a tissue framework, referred to as the canvas. In this model, the initial canvas consists of approximately 2,300 elements and measures 30 µm in both width and length. Each factor is assigned a value at either the segments or vertices of the canvas, where it promotes growth rates through a linear function (**Extended Data Fig. 8**). The model integrates two interconnected regulatory networks:

– The Polarity Regulatory Network (PRN), which defines tissue polarity and orients growth along the canvas.
– The Growth Rate Regulatory Network (GRRN), which modulates the local growth rates.

Polarity is established by the propagation of the factor POL, from a proximal organizer (PROXORG) toward a distal organizer called DISTORG. In the most realistic divergent model, DISTORG is expressed at the distal petal margin where it consumes POL, creating a divergence in the polarity field. This causes the growth direction to shift away from the petal midline and toward the margins in the distal region. Another key factor, DISGRAD, is predominantly expressed in the distal region, where it promotes growth perpendicular to the mid-petal axis (Kpar, or Kper). This model successfully replicates *Arabidopsis thaliana* petal development and is supported by molecular data, which, in accordance with our observations, suggest that *JAG* functions as a distal factor DGRAD and that auxin biosynthesis and signaling are involved in the establishment of the polarity field. Although this model accurately simulates *A. thaliana* petal growth, it fails to reproduce the petal shape of *Capsella* species, despite very similar overall growth rates (**Extended Data Fig. 9**). In *C. grandiflora*, petal growth dynamics are similar to those of *A. thaliana*, but the length-to-width ratio is significantly lower. Simply increasing Kper with the basic growth parameter did not generate realistic *Cg* petal shapes, as it only widened the distal region, maintaining a lobed shape, whereas *Cg* petals are generally more rounded. More realistic outlines were obtained by expanding both the DISTORG and DGRAD domains.

However, reducing DGRAD activity in this modified model did not significantly decrease petal length, as observed in *crjag* mutants (**Fig. 3**). To address this discrepancy, we modified the model so that DGRAD also affected Kpar, which resulted in petals highly similar in shape and size to those of *Cg*. Furthermore, abolishing DGRAD activity while accounting for the fragmented distal auxin distribution in *jag* petals (as in ^45^) led to a significant decrease in petal length, resulting in ‘jagged’ petals approximately 1.5 mm long, closely resembling *crjag* mutants. The models are provided as supplemental dataset and the parameters of the models used are presented in **Extended Data Table 7**.

Leaf growth was modeled using the organizer-based fixed polarity model described by Kuchen et al. (2012) ^46^. In this model, tissue polarity is established by diffusion of a polarity factor (POL) from a proximal organiser (PROXORG) at the base of the canvas. After an initial equilibration phase, the POL gradient is fixed and deforms passively with the tissue during growth. Growth parallel to the polarity field is promoted by PGRAD, while LAM promotes growth perpendicular to the polarity. MID inhibits perpendicular growth along the midline, and LATE modulates both parallel and perpendicular growth. All growth regulators remain fixed to their initial spatial distributions throughout development. Because crjag mutant leaves exhibit a 14% reduction in length and an 18% reduction in width, we assumed that JAG contributes similarly to both PGRAD and LAM activity. The model was therefore calibrated to reproduce the phenotypic differences observed in crjag leaves. We then ran both the petal and leaf models for the same number of iterations while varying the level of *JAG* activity according to its overall contribution to organ growth.

### Population genetics

We focused our analysis on polymorphisms segregating between the genotypes used for transgenic analysis, since they carry the alleles responsible for variation in petal size between mating strategies. Multiple sequence alignment of the four accessions’ JAG sequences (*C. rubella*, *C. grandiflora Cg1* and *Cg2*, and *C. orientalis*) was performed using MAFFT. Polymorphisms were identified from this alignment, which was then used to generate a consensus JAG sequence including all indels and the most frequent allele at each polymorphic site, using EMBOSS cons (parameters: -setcase 0-plurality 1). This consensus sequence replaced the native *JAG* sequence from the *C. rubella* reference genome (version 183).

We used this custom reference for read mapping of publicly available resequencing datasets (GATK pipeline), including 180 *C. grandiflora* individuals from a single population, 51 *C. rubella*, and 16 *C. orientalis* ^71–73^. The resulting VCF file contained 3,300 polymorphisms in the *JAG* coding and cis-regulatory regions. Indels were normalized using bcftools (norm followed by filter --IndelGap 5), applying parsimony and left-alignment, which reduced the set to 2,371 polymorphisms. We further curated the VCF applying the following filters: (i) remove polymorphisms located within large indels based on haplotypes used in transgenic experiments (accounted for ∼75% of the polymorphisms) (ii) remove polymorphisms absent from the genotypes used for introgression and complementation experiments; (iii) simplify indel annotations not fully resolved during normalization; and (iv) inspect ambiguous sites using IGV to resolve ambiguous or missing information. After filtering, the final dataset contained 340 polymorphisms for the Cg-Cr comparison and 536 for the Cg–Co comparison, among which 164 were shared. Among the 164 shared polymorphisms, i.e. the major allele in *Cg* differs from the major allele of *Cr* and *Co*, 84 were called convergent, i.e. the fixed allele in Cg (frequency >75%) differs from the fixed allele in Cr and Co (frequency >90%).

To facilitate downstream association analyses, multiallelic sites in the VCF were decomposed into biallelic pseudo-loci. For each multiallelic site, the list of alternative (ALT) alleles was extracted, and each ALT allele was treated as a separate locus. Genotypes were recoded as allele counts (0, 1, or 2 copies of the ALT allele), with missing data set to NA. This procedure was applied to all variants, and the resulting expanded genotype matrix was used for downstream analyses.

### Phenotypic association analysis and detection of overdominance

Phenotypic measurements for *Cg* individuals with available genomic information were retrieved from ^26^ and filtered to remove individuals with missing values. Genotypic data were converted into an individual-by-locus matrix, coded numerically, and merged with the phenotypic dataset. For each trait–locus combination, we tested for overall genotype effects using linear models with the genotype as a categorical predictor. Loci with fewer than 10 individuals in any genotype class were excluded from analysis. The overall significance of genotype effects was assessed by ANOVA, and per-term p-values were adjusted for multiple testing using the Benjamini–Hochberg false discovery rate (FDR) procedure.

To formally test for overdominance, we applied an additive–dominance model in which heterozygotes were explicitly coded as a binary variable. For each locus–trait pair with adequate sample size in all three genotype classes, we fitted the model Phenotype ∼ additive + dominance and identified significant dominance effects (p < 0.05) where heterozygotes exceeded both homozygotes in mean phenotype. Overdominant candidates were defined as loci that (i) showed a significant dominance term, and (ii) had heterozygous mean phenotypes greater than both homozygotes in the filtered dataset. Overlap between these statistical and phenotypic criteria was used to generate the final list of overdominant loci. Candidate loci were annotated with SNP metadata and visualized using boxplots and Manhattan-like genome position plots, with overdominant loci highlighted. The list of phenotype based overdominant polymorphisms was compared to the list of convergent polymorphisms.

### Identification of overdominant loci through inbreeding coefficient

Observed genotype counts per locus (*n*AA, *n*AB, *n*BB) were calculated from the expanded genotype matrix. From these, observed heterozygosity (*H*ₒ), expected heterozygosity (*H*ₑ), and the inbreeding coefficient (*F*_IS_= 1 − *H*ₒ/*H*ₑ) were computed. Hardy–Weinberg equilibrium (HWE) exact tests were performed using the HardyWeinberg package in R. Loci were classified as overdominant if they satisfied all of the following criteria: (i) significant deviation from HWE (*p* < 0.05), (ii) negative *F*_IS_ values (heterozygote excess), and (iii) *H*ₒ > *H*ₑ. Chromosome-wise plots of *F*_IS_ against genomic position were generated, marking overdominant loci and convergent overdominant (based on phenotype) candidates.

### Simulation of the influence of overdominant selection on allele trajectories

We conducted a common garden experiment at the University of Potsdam, Germany, using an F₂ population derived from a cross between *Cg* and *Cr*. Plants homozygous for the *Cg* (functional) allele at the self-incompatibility locus were selected, and their petal size was quantified. Reproductive efficiency was estimated as the number of seeds produced per fruit (which reflects animal pollination), normalised by the average number of ovules developed by each individual and used as a proxy for pollination-mediated female fitness.

The relationship between petal size and reproductive efficiency was assessed using Pearson’s correlation. A linear regression model was then fitted with reproductive efficiency as the response variable and petal area as the predictor. Regression coefficients and R² values were extracted to describe the strength and direction of the relationship. For visualization, scatterplots with fitted regression lines and annotated regression equations were generated.

Allele frequency dynamics were simulated using a discrete-generation Wright–Fisher model. The model incorporated: Effective population size *N_e_* =500,000, 10^6^ generations and an initial allele frequency *p_0_* =0.1 and selection gradient derived from genotype-phenotype relationships. For each replicate (500 replicates), genotype frequencies were adjusted according to selection coefficients, and the next generation was sampled multinomially to account for genetic drift. Simulations were also conducted under neutrality (no selection) for comparison.

Fitness of genotypes was modelled linearly as a function of deviation from the mean phenotypic value:

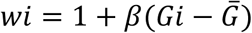

where *β* is the selection coefficient relating phenotype to fitness, *Gi* is the genotype’s phenotypic value, and *G*^-^ is the population mean. Overdominance was assessed by comparing heterozygote fitness to the two homozygotes. A stable internal equilibrium allele frequency *p*^ was estimated when overdominance was detected.

For each simulation replicate, we recorded allele frequencies across generations and at the final generation: homozygote and heterozygote counts, observed heterozygosity (*Ho*), expected heterozygosity (*He*), and *Fis*. Hardy–Weinberg equilibrium was tested using exact tests, and the proportion of replicates deviating from equilibrium at α=0.05 was reported.

Allele frequency trajectories were summarized as mean ± 95% confidence intervals across replicates. Final-generation allele frequencies were visualized as histograms, and probabilities of maintaining polymorphism were estimated as the proportion of replicates with 0 < heterozygote frequency < 1.

### Statistical analyses

Statistical analyses were performed using R (version 4.2.1). Two-tailed Student’s t-test assuming unequal variances was used for two-sample comparisons. For multiple comparisons, we conducted Tukey’s HSD tests, performed using the R/agricolae package. Data visualization was done using the R/ggplot2 package or R/base. In all boxplots, the middle lines represent the median, the lower and upper hinges correspond to the first and third quartile, respectively. The upper and lower whiskers add or subtract 1.5 interquartile ranges to/from the 75 and 25 percentile, respectively. Overdomiance was tested using Linear mixed-effects models,fitted using the lmer function in the lme4 package. Estimated marginal means were obtained using the emmeans package. Planned contrasts between genotypes were tested with Bonferroni-adjusted p-values, and degrees of freedom were approximated using the Kenward-Roger method.

## Data availability

The genome sequence data used in this manuscript have been previous deposited to NCBI’s Sequence Read Archive (SRA) or the European Nucleotide Archive (ENA) under the accession number SRA: PRJNA1233196, PRJNA1233196, PRJNA275635 and PRJEB6689 and are publicly available. All materials used in this study are available upon request.

## Acknowledgments

We thank Fredric Hedlund and Kathrin Hesse for plant care and members of the Lenhard and Rosa groups for discussion and comments on the article. The computation and data handling were provided by the Swedish National Infrastructure for Computing (SNIC) at Uppmax, partially funded by the Swedish Research Council through grant agreement no 2018-05973. This work was supported by: Deutsche Forschungsgemeinschaft grant SI1967/2 (AS) Swedish Research Council grant 2018-04214 (AS, KS) and theNovo Nordisk Fonden NNF22OC0079830 (AS, KS)

## Author contributions

K.S., N.W. and A.P. performed and analysed the experiments. U.F. generated transgenic lines, and T.F.L. contributed to phenotypic analyses of mutant lines. C.K. carried out the bioinformatic analyses. A.S. acquired funding, administered the project and, together with M.L., S.R. and Y.D., supervised the research. A.S., N.W., K.S. and A.P. drafted the manuscript, and all authors contributed to revising and editing the final version.

## Ethics declarations Competing interests

The authors declare no competing interests.

